# Immigration rates during population density reduction in a coral reef fish

**DOI:** 10.1101/033274

**Authors:** Katrine Turgeon, Donald L. Kramer

## Abstract

Although the importance of density-dependent dispersal has been recognized in theory, few empirical studies have examined how immigration changes over a wide range of densities. In a replicated experiment using a novel approach allowing within-site comparison, we examined changes in immigration rate following the gradual removal of territorial damselfish from a limited area within a much larger patch of continuous habitat. In all sites, immigration occurred at intermediate densities but did not occur before the start of removals and only rarely as density approached zero. In the combined data and in 5 of 7 sites, the number of immigrants was a hump-shaped function of density. This is the first experimental evidence for hump-shaped, density-dependent immigration. This pattern may be more widespread than previously recognized because studies over more limited density ranges have identified positive density dependence at low densities and negative density dependence at high densities. Positive density dependence at low density can arise from limits to the number of potential immigrants and from behavioral preferences for settling near conspecifics. Negative density dependence at high density can arise from competition for resources, especially high quality territories. The potential for non-linear effects of local density on immigration needs to be recognized for robust predictions of conservation reserve function, harvest impacts, pest control, and the dynamics of fragmented populations.

## Introduction

The connection between density and dispersal has important implications for ecology, evolution, and behavior [1–5]. Over long distances, density-dependent dispersal can influence range expansion and the extinction and colonization of patches [4,6]. When dispersal involves localized movements within populations, density dependence can affect other density-dependent processes such as reproduction, growth, and mortality. In addition, it can affect source-sink dynamics, the function of conservation reserves and reintroduction programs, and the control of pests and invasive species, for example, compensating for localized mortality and increasing movements of animals from protected to harvested areas [7–12].

Numerous empirical studies provide evidence for an association between emigration rate and population density [13,14]. However, the broader relationship between local density and dispersal is not well understood, in part because both theoretical and empirical investigations have tended to focus on emigration. The effects of density on the equally important movement and immigration stages of the dispersal process have been largely ignored [5,13,15]. The substantial literature examining the influence of conspecifics on the number of individual marine fish and invertebrate larvae that enter local populations from the plankton [16–18] appears to be an exception. However, this literature has only limited application to questions of density-dependent immigration because the number of individuals arriving (called 'settlement' in this literature) often cannot be distinguished from the number surviving to some early stage ('recruitment'), which itself is often density-dependent [18].

Immigration may increase as density increases if Allee effects [19] occur or if conspecific abundance provides an indication of habitat quality [20]. On the other hand, immigration may decrease as density increases because higher densities result in lower *per capita* resource availability or lower quality of locations available for settlement [21]. Since the benefits of increasing density are more likely when densities are relatively low and the costs of increasing density are more likely when densities are relatively high, a hump-shaped relationship between immigration and density may be expected when density varies over a broad range [22]. Alternatively, density might have no influence on immigration (*i.e.* density independence) if animals are unable to assess density or resource availability, if populations never approach carrying capacity, if Allee effects are unimportant, or if immigration rate is limited by the availability of potential immigrants.

Empirical evidence to indicate whether immigration rate shows a hump-shaped response to density is sparse. Only a few field studies have examined immigration rate over a sufficient number and range of density levels to detect a hump-shaped pattern, if one were present [22–30]. Some studies have presented immigration rate data only after dividing by the number of residents, potentially obscuring lower immigration to sites with fewer residents (discussed below). Finally, almost no studies have used a replicated, experimental design to control for potentially confounding correlates of density and immigration such as habitat quality, functional connectivity, and immigrant availability [22,24,27].

In the present study, we examine whether adults of two small, interspecifically territorial coral reef damselfishes show density-related variation in local immigration in response to the experimental manipulation of density from saturation to complete absence of conspecifics. On 7 sites surrounded by suitable, occupied habitat, we repeatedly removed a small, constant proportion of fish, continuing until no fish remained and immigration ceased. Previously, using data from this experiment, we showed that the total number of immigrants arriving over the entire experimental period was influenced by inter-site differences in the availability of potential immigrants in adjacent areas and by the relative habitat quality between the depletion and source areas but not by differences in landscape connectivity [31]. Two control sites had almost no immigration during equivalent observation periods. Here, we examine the number of immigrants arriving after each removal event to ask how immigration rate within sites changes in response to the gradual reduction of density.

## Methods

### STUDY SPECIES

Adult and subadult longfin *Stegastes diencaeus* and dusky damselfish *S. adustus* (formerly *S. dorsopunicans),* hereafter ‘damselfish’, of both sexes defend small (about 1 m^2^) territories on dead coral substrate. These territories provide food (benthic algae), shelter holes, and nest sites for males [31,32], and there are no non-territorial ‘floaters’. Because they are diurnal, active, conspicuous, and occur at high density, they are well suited for studies of immigration. Their fringing reef habitat is a naturally fragmented landscape in which continuous reef and patches of various sizes are separated by a matrix of sand and rubble. Damselfish seldom move far from solid reef substrate, so sand and rubble patches form partial barriers to movement within a reef [33]. Dispersal occurs at two life history stages and spatial scales. After hatching from eggs guarded by males, larvae disperse in the plankton, later settling on the reef as small juveniles. Thus, individuals are unlikely to be closely related to their neighbors, and genetic differentiation is unlikely over relatively large spatial scales [34]. Upon reaching adult size, mortality rate is low, and spontaneous moves are rare [35]. Nevertheless, damselfish relocate quickly in response to vacancies in higher quality habitat [35,36]. This local dispersal or ‘home range relocation’ of adult and subadult individuals, almost certainly from within the same fringing reef and likely from relatively close territories, is the focus of our study. It is similar to the within-patch dispersal of many mammals and birds [30,37,38]. In our study area, adult territories are contiguous on suitable substrate, with their number and spatial extent relatively stable over the annual cycle as well as between years (K. Turgeon, *unpublished data*), as also noted for other territorial damselfishes [39,40]. Stable densities and distributions suggest that these populations are close to saturation on fringing reefs in Barbados.

### REMOVAL EXPERIMENT

Our study design mimicked a situation in which localized predation or human exploitation created an attractive sink for immigrants from surrounding source areas that were not subject to the same mortality. To the best of our knowledge, this approach has not been used in previous studies of density-dependent immigration. Although repeated removals may reduce the number of potential immigrants, examining the density-immigration relationship during gradual depletion provides a rare opportunity to examine immigration over a broad range of densities while holding other ecological features constant, as well as providing insight into the impacts of localized harvest and pest removal on dispersal.

We conducted the experiment on seven experimental and two control sites in the spur and groove zones of five fringing reefs [experimental: Bachelor Hall (1), Heron Bay (4), Sandy Lane (2); control: North Bellairs (1), South Bellairs (1), S1 Fig.] along the west coast of Barbados (13°10’N, 59°38’W). Each site consisted of a rectangular depletion area of about 150 m^2^ within the much larger spur and groove zone of the fringing reef. We defined a 10-m wide band around the depletion area as a source area for potential immigrants, based on the distances of extraterritorial forays which could serve to detect vacant territories [31,41,42]. We selected experimental sites so that the depletion areas were similar in number of residents, total area, and area of hard substrate (S1 Table). Using natural variation in the shapes of the reefs, we selected source areas to vary in abundance of potential immigrants, quality of the habitat and landscape connectivity [S1 Table; see Fig. S1 in [31] for examples].

In the depletion area at each experimental site, we marked all resident damselfish > 5 cm (total length) with visible implant elastomer (Northwest Marine Technology, Shaw Island WA, USA) and mapped their territories. Experimental sites were observed at 2 or 3-day intervals for two weeks prior to removal. Then, every 2 or 3 days, we removed a constant number of randomly selected individuals, including both residents and the immigrants that had replaced previously removed residents, equal to about 15% of the original population (i.e., 7 - 10 individuals per event, depending on the number of residents). In one case (site HB1), the sequence was delayed for several days by a hurricane. The day after each removal, we mapped all occupied territories in the depletion area, noting the presence of immigrants (untagged individuals). Approximately 30 hours was considered sufficient to achieve a new equilibrium because about 90% of recolonisations in both species occurred by the end of observations on the second day. Previous studies also found that territory recolonisation in damselfishes often occurs within one day [35,36,43]. Removals continued until no fish remained and immigration had ceased for at least 6 days. (In 2 sites, a single, small individual could not be captured.) Complete trials required approximately five weeks, depending on immigration rate. In the source area, we counted all damselfish > 5 cm once but for logistical reasons were not able to monitor changes in density or position at each removal event. Because of the intensity of data collection, we were able to conduct only one trial at a time. Trials took place between April and September 2005-2007.

### ETHIC STATEMENT

Fish removal was needed to determine if movement occurs when the density is reduced. Prior to the experiment, specimens were marked with an injection of a small amount of inert elastomer under the scales (VIE tag - Northwest Marine Technologies). The tagging procedures required less than 30 s to perform and mortality from this method of tagging is extremely low. To reduce fish stress of these two highly territorial species, we marked them underwater, without anaesthesia, near the site of capture so that they can return to their territories quickly. During the removal experiment, fish were removed using cast nets whenever possible. When cast nets were not successful, we used micro-spearfishing. After capture, fish were immediately euthanized using an overdose of carbon dioxide prepared by dissolving antacid powder (Eno, approx. 5g/L) in sea water. The effects of this localized fish removal on the population and community are similar to the harvest effect produced by fisheries and were expected to have modest effects on competition and predation. Our research complied with the ABS/ASAB Guidelines for the Treatment of Animals in Behavioral Research and Teaching and was approved by the McGill University Animal Care Committee, Animal Use Protocol 5039. Field work did not involve endangered or protected species. No specific permissions were required to harvest these two damselfish species in Barbados.

### DATA ANALYSIS

#### Scaling immigration rate

We carried out separate analyses of immigration rate using per site scaling (total number of immigrants arriving on the site following each removal event), per new vacancy scaling (number of immigrants/number of damselfish removed at each event), and *per capita* scaling (number of immigrants/number of damselfish present following each removal event). Per site scaling provides the most direct measure of immigrant movement and is equivalent to per unit area scaling. We included per new vacancy scaling because territory availability was the limiting resource and relating immigration to vacancies was therefore likely to illuminate the processes involved [*e.g*., [25]]. Because 92% of all immigrants (n = 122) settled on territories from which the owner had been removed during the immediately preceding event rather than during previous removal events, we scaled to the number of new vacancies rather than total vacancies. Even though removal of a constant number of damselfish at each event means that per site and per new vacancy scaling would produce similar patterns over most of the density range, important variation could occur at low densities where the total number of remaining damselfish was less than the number to be removed. *Per capita* scaling was included for completeness because demographic rates, including emigration and immigration, are normally scaled to the population size in analyses of density dependence [44]. However, it is important to recognize that relating immigrants per damselfish to number of damselfish is an example of spurious correlation (Y/X vs. X) that can produce a negative correlation even if immigration rate per site does not change with density [45]. This is referred to as ‘apparent’ or ‘pseudo-density dependence’ [18]. For per site scaling, we included data on immigration before the start of removal as well as data after the number of residents had been reduced to zero. Per vacancy scaling could not include data before any fish had been removed and *per capita* scaling could not include data after density had been reduced to zero, because the values were undefined.

#### Statistical analyses

To assess density dependence of immigration, we used curve-fitting analyses to compare the fit of 6 alternative functions for each site and then examined the generality of the pattern with Generalized Additive Mixed effect Models (GAMM) on the combined dataset. Both analyses examined the relationship between the total number of damselfish still present on each site after each removal event, including both residents and earlier immigrants (independent variable), and the number of immigrants arriving at the site between that removal and the subsequent count (dependent variable). For the independent variable, number of damselfish was equivalent to density because area was constant within sites. For the dependent variable, we scaled the number of immigrants in three ways to facilitate interpretation as described above. We combined both species for this analysis because territories were defended interspecifically, because there was too much variation in relative numbers of the two species between sites and between removal events for separate analyses by species, and because a preliminary analysis did not indicate an effect of species on the response to density (K. Turgeon, *unpublished analysis*).

For each site, we compared the fit of six alternative functions corresponding to density independence, linear negative or positive density dependence, exponential negative or positive density dependence, saturating positive density dependence, and two functions that can model a hump-shaped combination of negative and positive density dependence (Table 1). Including alternative functions (linear and exponential for monotonic functions, polynomial and Ricker for hump-shaped relationships) allowed greater flexibility in the fit. The saturating positive density-dependent function addressed the potential for immigration to be constant and then drop at lower density as the source became depleted. Parameters for each function were estimated with maximum likelihood approach with the mle2 function (normal probability distribution) available on the bbmle package version 1.0.4.1 [46]. To compare the support of each function to explain observed immigration, we used the Information Theoretic Approach [47]. We used Akaike's Information Criterion modified for small sample sizes (AICc) to assess the fit of each function to observed data. Lower AICc values indicated a better fit. We calculated the difference between AICc for each model *i* and the lowest observed AICc (ΔAICc) and compiled normalized Akaike weights (*w*_*i*_) the probability that function *i* is the best function, given the data and set of candidate functions [48]. For each site, we also computed the “evidence ratio” to compare the relative likelihood of any two functions (*w*_*i*_/*w*_*j*_; [48]). To determine the best function overall, we calculated the mean Akaike weights (mean *w*_*i*_) for each function.

**Table 1.** Alternative functions fit to the data relating immigration (*I*_*t*_) to the number of damselfish remaining on the site (*D*_*t*_) following each removal event, where *a* and *b* are estimated parameters. For per site scaling, *I*_*t*_ equaled the total number of immigrants arriving on the site between the removal and the subsequent count. For per new vacancy scaling, *I*_*t*_ equaled the total number of immigrants arriving on the site between the removal and subsequent count divided by the number of damselfish removed. For *per capita* scaling, *I*_*t*_ equaled the total number of immigrants arriving on the site between the removal and the subsequent count divided by the number of damselfish remaining immediately after the removal.

| Function | Pattern |
| --- | --- |
| $I_t = b$ | 1) Density independent |
| $I_t = aD_t + b$ | 2) Linear negative or positive density dependence |
| $I_t = ae^{-bD_t}$ | 3) Exponential negative or positive density dependence |
| $I_t = a(1 - e^{-bD_t})$ | 4) Saturating positive density dependence |
| $I_t = -aD_t^2 + bD_t$ | 5) Quadratic polynomial (hump-shaped) |
| $I_t = aD_te^{-bD_t}$ | 6) Ricker (hump-shaped) |

For the combined analysis, we used Generalized Additive Mixed effects Models (GAMMs) because they combine the utilities of linear mixed models [49] and Generalized Additive Models (GAM; [50] so that random factors, fixed factors and nonlinear predictor variables can all be estimated in the same model. Rather than fitting curves as in the site analysis, GAMMs model nonlinear relationships between density and immigration as a smoothing function. Because of the potential temporal autocorrelation between successive removal events and the risk of violating the assumption of residual independence in interpreting *p*-values, we used an autoregressive correlation structure (corAR1) in the GAMM. Site identity was used as a random factor, and the autocorrelation in error structure was accounted for by nesting the removal event sequence within site. To make sites more comparable, the independent variable was changed from number of damselfish at each removal event to the proportion of the initial number. GAMMs for all three scales were fit using the mgcv package v. 1.7-28 in R [51,52].

## Results

Immigration occurred almost exclusively at intermediate densities. It never occurred during the pre-removal period when densities were maximal, but started after the first removal in three sites (SL1, SL2, BH1) and after the second removal in three others (HB3, HB2 and HB4) but only after the sixth removal in HB1 (Fig. 1). After the first occurrence of immigration in each site, immigration followed most subsequent removals until density became very low. In 6 sites, there was no immigration after density was reduced to 0 or to a single uncapturable individual. In 1 site (HB4), 2 individuals arrived after density was reduced to zero the first time. However, most sites did have 1 - 4 immigrants when density was first reduced to 1 or 2 individuals.

**Figure 1.**
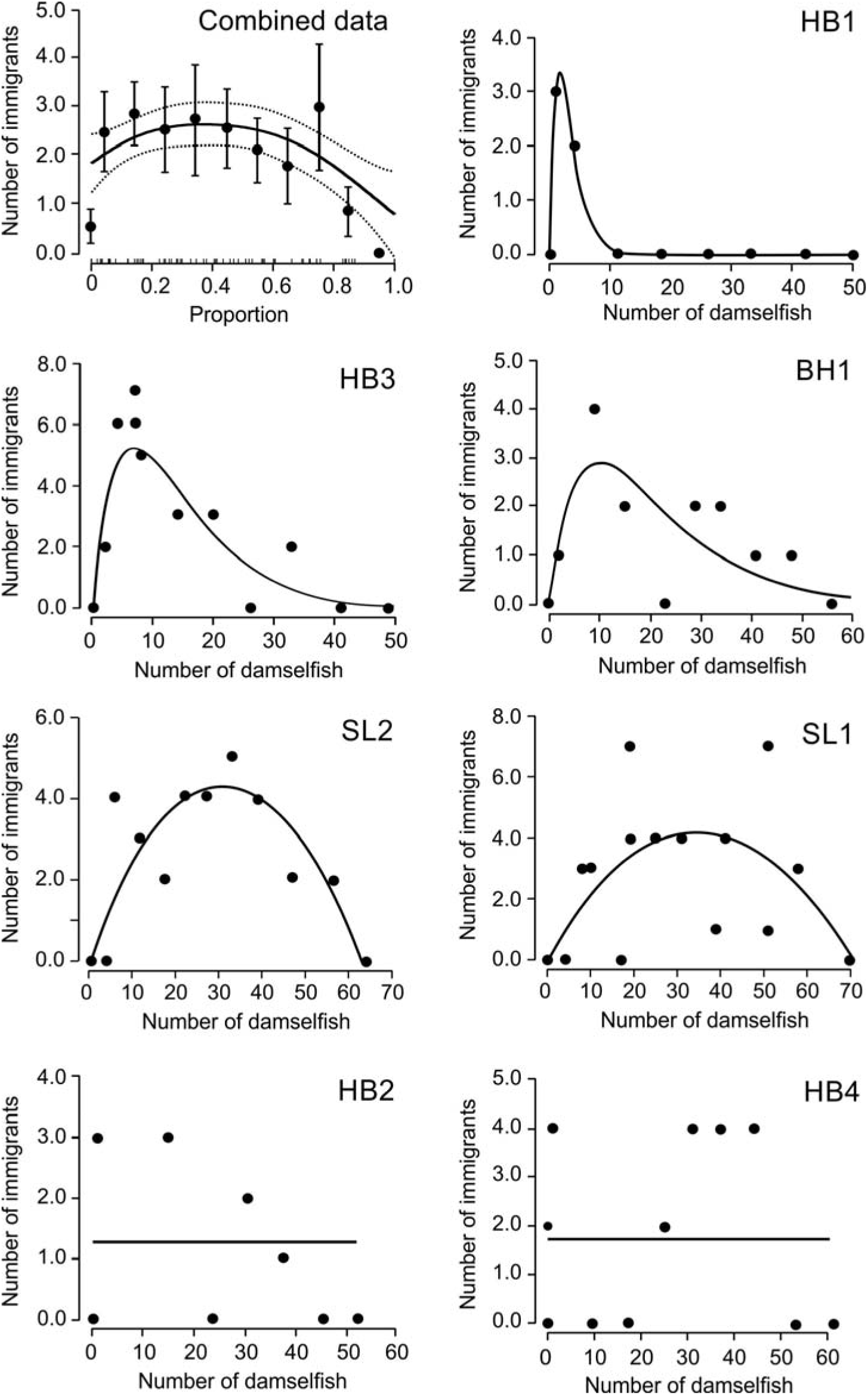
Relationship between the number of immigrants arriving on a site after a removal event and the number of damselfish remaining following each removal event (site scaling) for each individual site and for all sites combined (upper left panel). Individual sites are ordered by strength of support for a hump-shaped (Ricker or polynomial) function and identified by abbreviations (Bachelor Hall Reef: BH1; Heron Bay Reef: HB1, HB2, HB3, HB4; Sandy Lane Reef: SL1, SL2). Lines represent the best fitting empirical function based on maximum likelihood estimates (MLE) of parameter values. For all sites combined, the abscissa indicates the proportion of the initial number of fish on the site before removal. Each point represents the mean ± 1 SE of all points falling within bins of 0.1, with density = 0 shown separately. Sample sizes per bin vary depending on replacement rates. The short inside tick marks on the abscissa represent observed proportion values. The black line in the upper left panel represents the significant smoothing term function from a GAMM model and the dashed lines represent the Bayesian credible interval as calculated by the mgcv package v. 1.7-28 in R.

For per site scaling (*i.e.* the total number of immigrants arriving after each removal event), the curve-fitting analysis of individual sites indicated that the best function was hump-shaped density dependence in 5 sites (Ricker function in 3, polynomial in 2) and density independence in 2 sites (Table 2, Fig. 1). In comparison to density independence, the evidence ratio for density dependence was high (> 56) for 3 sites and moderate (2.6 – 3.0) in the other two. On average, the hump-shaped Ricker function provided the best fit as indicated by the mean *w*_*i*_ (Table 2), with a moderate evidence ratio compared to density independence (2.3). The fitted functions indicated that, on average, immigration per site continued to rise until density was less than half the starting density (peaks occurred between 6% – 50% of the starting population, mean = 29%, Fig. 1). For the combined dataset analysis, the GAMM supported a hump-shaped pattern (significance of the smoothing term; p = 0.014; Fig. 1). As density decreased, mean immigration rose to a broad peak between about 10 and 40% of the starting population, dropping to a very low level only when density approached zero. These patterns were strongly influenced by but not completely dependent on the very low immigration at zero density; when zero density data were excluded from the analysis, the hump-shaped pattern remained very strong in the GAMM analysis of the combined data and was most likely in 3 individual sites, but density independence became slightly more likely than hump-shaped density dependence in 2 sites for which a hump-shaped function fit best with zero density included (K. Turgeon, unpublished analyses).

**Table 2.**
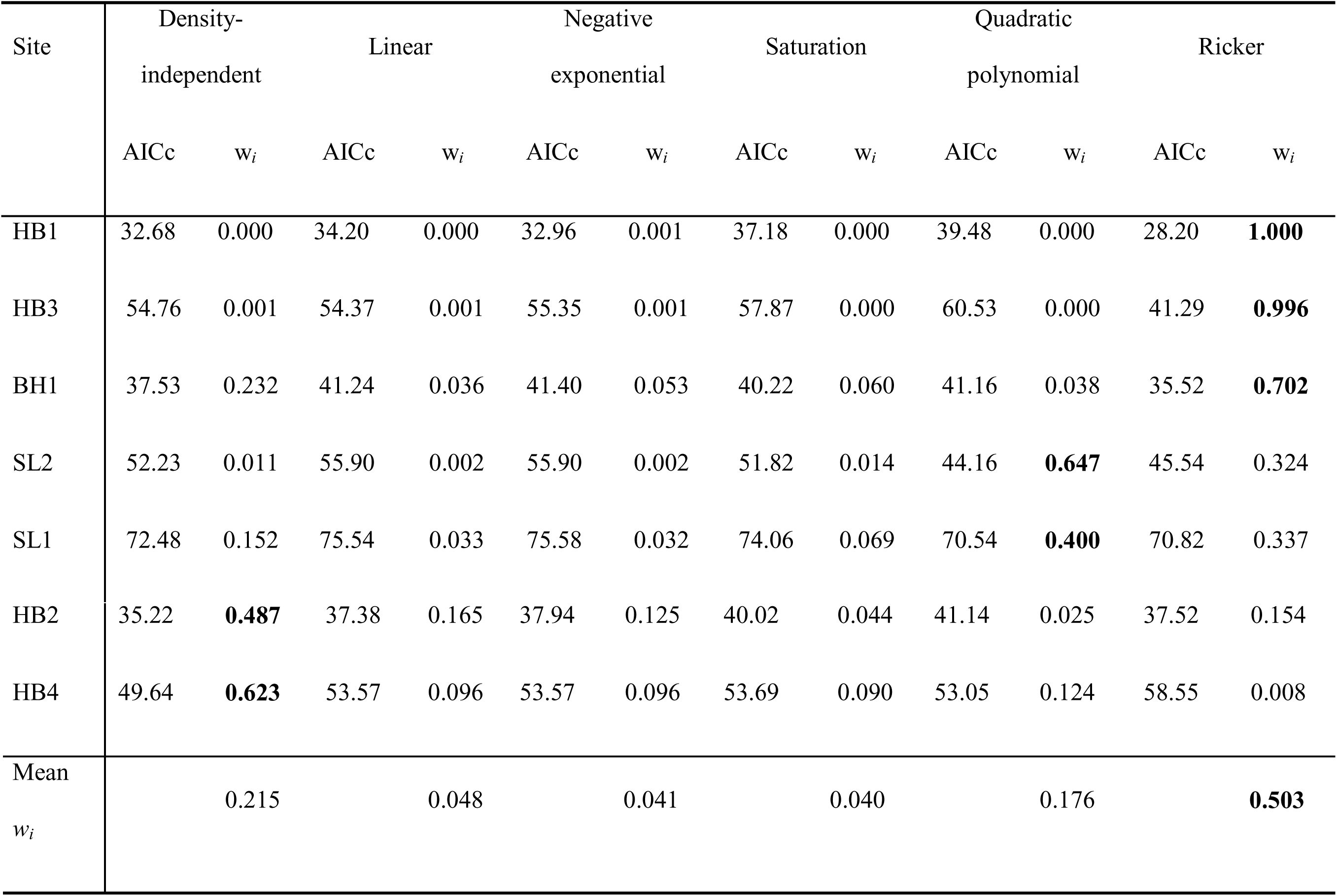
AICc scores (AICc) and Akaike weights (w_*i*_) for each of the six empirical functions predicting 289 the observed number immigrants arriving on the site following a removal event in relation to population size of damselfish following that event (site scaling) on seven experimental sites. The mean Akaike weight (Mean w_*i*_) for each function is given at the bottom each column. For each site and the mean of all sites, the function with the highest support is indicated by bold w_*i*_. Sites ordered as in Fig. 1. See Figure 1 for site abbreviations and Table 1 for the functions.

| Site | Density-independent |  | Linear |  | Negative exponential |  | Saturation |  | Quadratic polynomial |  | Ricker |  |
| --- | --- | --- | --- | --- | --- | --- | --- | --- | --- | --- | --- | --- |
| | AICc | $w_i$ | AICc | $w_i$ | AICc | $w_i$ | AICc | $w_i$ | AICc | $w_i$ | AICc | $w_i$ |
| HB1 | 32.68 | 0.000 | 34.20 | 0.000 | 32.96 | 0.001 | 37.18 | 0.000 | 39.48 | 0.000 | 28.20 | **1.000** |
| HB3 | 54.76 | 0.001 | 54.37 | 0.001 | 55.35 | 0.001 | 57.87 | 0.000 | 60.53 | 0.000 | 41.29 | **0.996** |
| BH1 | 37.53 | 0.232 | 41.24 | 0.036 | 41.40 | 0.053 | 40.22 | 0.060 | 41.16 | 0.038 | 35.52 | **0.702** |
| SL2 | 52.23 | 0.011 | 55.90 | 0.002 | 55.90 | 0.002 | 51.82 | 0.014 | 44.16 | **0.647** | 45.54 | 0.324 |
| SL1 | 72.48 | 0.152 | 75.54 | 0.033 | 75.58 | 0.032 | 74.06 | 0.069 | 70.54 | **0.400** | 70.82 | 0.337 |

Table 2. Continued and concluded
| Site | Density-independent |  | Linear |  | Negative exponential |  | Saturation |  | Quadratic polynomial |  | Ricker |  |
| --- | --- | --- | --- | --- | --- | --- | --- | --- | --- | --- | --- | --- |
| | AICc | $w_i$ | AICc | $w_i$ | AICc | $w_i$ | AICc | $w_i$ | AICc | $w_i$ | AICc | $w_i$ |
| HB2 | 35.22 | **0.487** | 37.38 | 0.165 | 37.94 | 0.125 | 40.02 | 0.044 | 41.14 | 0.025 | 37.52 | 0.154 |
| HB4 | 49.64 | **0.623** | 53.57 | 0.096 | 53.57 | 0.096 | 53.69 | 0.090 | 53.05 | 0.124 | 58.55 | 0.008 |
| Mean<br>$w_i$ | | 0.215 | | 0.048 | | 0.041 | | 0.040 | | 0.176 | | **0.503** |

For per new vacancy scaling, the best function for individual sites was hump-shaped (Ricker function) in all 5 sites that showed a hump-shaped response in per site scaling and density-independent in the other 2 sites (S2 Table, S2 Fig. black dots and lines). Compared to density independence, the evidence ratio for density dependence was high in 3 sites (> 10.9) but low at the other 2 (1.3 – 1.5). On average, the Ricker function provided the best fit (Mean *w*_*i*_; S2 Table) but was only slightly more likely than density independence (evidence ratio = 1.6). The proportion of new vacancies occupied by immigrants following individual removal events ranged from 0 to 1 but was almost always less than 0.3 (median = 0.25, 95% CI = 0.20 – 0.31, N = 77; S2 Fig. black dots). For the combined dataset analysis, the GAMM supported a hump-shaped relationship between the number of immigrants per new vacancy and the proportion of the initial abundance (significance of the smoothing term; *p* = 0.016; S2 Fig.).

For *per capita* scaling, the best-fitting function was density-dependent for all 7 sites (S3 Table, S2 Fig. grey lines and dots). The best function was hump-shaped (Ricker) for the 5 sites that showed hump-shaped relationships with per site and per new vacancy scaling, negative exponential for 1 site that had been density independent with per site and per new vacancy scaling and tied between hump-shaped (Ricker) and negative exponential for the other. The evidence ratio for density dependence (Ricker and negative exponential) compared to density independence was very high (> 100) for 5 sites and moderate and high for the other 2 (2.1, 6.7). On average, the best fit was hump-shaped (Ricker; Mean *w*_*i*_; S3 Table) with high evidence ratios in comparison to density independence (16.8) and the negative exponential (6.1). For the combined dataset analysis, the GAMM supported a negative relationship between the number of immigrants *per capita* and the proportion of the initial abundance (significance of the smoothing term; *p* < 0.001; S2 Fig.).

## Discussion

Patterns of damselfish immigration in response to repeated small removals provided evidence that immigration rate was non-linearly density-dependent with a maximum at intermediate density. This relationship was supported by the GAMM analysis of the combined data, by the presence of immigration at intermediate densities in contrast to the near absence of immigration at the lowest and highest densities at all sites, and by support for a hump-shaped relationship in 5 of 7 individual sites as well as the Akaike weights averaged over all 7 sites. The statistical patterns for individual sites were similar with both site and per new vacancy scaling. This was expected because the number of fish removed at each event was constant for each site over most of the trial. However, although per new vacancy scaling did not allow inclusion of the starting density before the first removal, it did provide additional support by correcting for the limited number of new vacancies generated by removal events at the lowest densities when the number of remaining individuals was less than the prescribed number of individuals to be removed.

There was less support for density independence than for hump-shaped density dependence. Density independence, the null hypothesis, was the best fitting model in 2 sites, and a good second alternative to hump-shaped functions in two others and in the mean Akaike weights. Note that support for density independence was based not on constant immigration over densities but on variable immigration that was not fit better by any alternative pattern. Because there was only a single trial at each site, we cannot be sure that there are true differences in the pattern of density dependence among sites. We found no consistent relationships between site characteristics and strength of support for a hump-shaped function (K. Turgeon, unpublished analyses). Although other studies have identified spatial variation in immigration rates [24,29], we are not aware of any studies that have examined differences among sites in the patterns of density dependence. Variable patterns in territory availability arising from random selection of damselfish to be removed could explain some of the variation in the fit of alternative functions. For example, removal events in which more of the vacated territories were of high quality, closer to abundant potential source populations, or adjacent to each other early in the sequence would have been likely to favor higher immigration at high densities, thereby reducing support for a hump-shaped pattern. Conversely, a high proportion of poor-quality territories made available in the middle part of the sequence would reduce immigration where high immigration was expected. Whether due to differences in factors influencing density dependence or to the stochastic effects of removal patterns or both, among-site variation in the pattern of density dependence highlights the importance of replication in such dispersal studies, despite the logistical challenges of achieving it.

The support for negative density dependence with *per capita* scaling was likely a mathematical artifact. In the two sites in which the density independent function provided the best fit with per site scaling, there was a good fit to the negative exponential with *per capita* scaling. Because a negative statistical trend is the expected result of relating Y/X to X when Y shows no relationship to X, this is probably a spurious correlation, sometimes referred to as pseudo-density dependence [18,45]. Apparently, the hump-shaped pattern was sufficiently strong that it still provided the best fit with *per capita* scaling in the other 5 sites and in the average for all sites (S3 Table). However, the stronger negative trend resulting from *per capita* scaling is apparent in all sites (S2 Fig.), resulted in increased support for the negative exponential in 2 sites and the average for all sites (S3 Table), and is reflected in the GAMM on the combined data, which indicated a simple negative relationship. This pattern also may have been affected by limitations of the GAMM for modeling the extremely sharp changes. [18], reviewing cases of larval fish recruitment to coral reefs, noted that diverse patterns of positive, negative or neutral density dependence with site scaling all became negative with *per capita* scaling. [30] also found that positively density-dependent immigration of red-backed shrikes to their population became negative with *per capita* scaling. Unlike other demographic variables, the numerator and denominator of *per capita* immigration are derived from different populations, necessitating that patterns be interpreted with caution.

Our study provides the first experimental evidence for a hump-shaped relationship between immigration rate and local density. Only three observational studies have also found hump-shaped relationships. Spiny lobsters (*Panulirus argus*) immigrating into marine reserves when disturbed by sport fishing showed a hump-shaped relationship between density in the reserves and immigration per site [28]. Hump-shaped relationships in larval settlement, probably not confounded by post-settlement mortality, have also been reported in barnacles *Semibalanus balanoides* [23] and humbug damselfish *Dascyllus trimaculatus* [24].

A positive relationship between density and immigration similar to the one we found between low and intermediate densities has been reported in several previous studies. In studies covering a broad density range, density and immigration per site or per vacancy were positively associated in three studies of birds [25,26,30] and in two studies of larval crabs and fishes [22,27]. Studies with a more limited range of densities (often simply presence/absence) also provide evidence for a positive effect of conspecifics in birds (where it is referred to as 'conspecific attraction' and often studied using models and song playback rather than established conspecifics, reviewed by [53], in marine invertebrate larvae (where it is referred to as 'gregarious settlement' reviewed by [16], and in *Dascyllus* damselfish larvae [54–56].

Removal experiments on a variety of territorial species suggest that a negative relationship between density and immigration like that we found between intermediate and high densities should be common, but there is relatively little support for this relationship in studies covering a broad density range. Studies designed to test for the presence of a surplus breeding population in territorial birds have often found rapid, complete or partial replacement of removed territory holders [reviewed by [57]]. Similar patterns have been found in some mammals [*e.g.,* [10]], reptiles [*e.g.,* [58]], fishes, including territorial damselfishes (*e.g.,* [59,60]], and insects (*e.g.,* [61]]. This suggests that immigration might be proportional to the reduction in density, but direct evidence over a range of densities is limited. An experiment using small juvenile humbug damselfish showed a nonlinear negative relationship [24], although *per capita* scaling makes interpretation difficult. An observational study on song sparrows *Melospiza melodia* immigrating to small islands reported a negative relationship in males but no relationship in females [29]. Two studies of larval fish recruitment covering a broad range of densities found a negative effect of adult density, but the study designs were unable to rule out post-settlement mortality as a factor [62,63]. Among studies with a limited density range, we are aware of only one showing a negative effect of density on larval fish settlement [64]. Thus, the negative trend we observed over most of the density range is only the third example of such a pattern over a broad density range. However, taken together, the literature suggests that both positive and negative trends may occur, suggesting that hump-shaped relationships may be observed more commonly if a sufficiently broad range of densities is studied.

The extremely low rate of immigration at very low densities in our study could be a consequence of vacant habitat avoidance (the inverse of conspecific attraction), reduced availability of potential immigrants, or both. If conspecific attraction exerts a powerful influence on habitat selection, individuals might avoid otherwise suitable habitat lacking conspecifics. Conspecific attraction can be favored by selection if the presence or abundance of other individuals indicates higher habitat quality (habitat cueing) or if higher density increases fitness (Allee effect) through improved predator avoidance, increased access to mates, or other benefits [20,65]. Previous studies that found positive correlations between density and immigration per site in nesting birds, spiny lobsters, as well as the settlement and recruitment of marine larvae have attributed the patterns to this phenomenon [22,25,26,28,66]. Allee effects might occur in damselfish because individuals with neighbors are more able to resist intrusions by competing herbivorous parrotfish and surgeonfish [60,67,68]. In addition, isolated damselfish have increased costs of mating and decreased choice because of the risks of longer excursions from their territory, especially if they have to travel over open sand [41,69].

An alternative explanation for very low immigration at the lowest densities is depletion of the supply of potential immigrants. Successive removals in the depletion area could reduce the pool of potential immigrants both by lowering the number of fish in the surrounding source areas and by providing more choice of territories for those individuals remaining. If immigrants moving to the depletion area were themselves quickly replaced by others from more distant areas, immigrant depletion effect would be minor, but if individuals were rarely replaced, the consequences could be substantial. Although we were not able to census the potential source populations after each removal during the experiment, we observed that many damselfish remained within 10-m of the borders of the depletion area even after we had removed the last individual. It is possible that these individuals remaining in the source areas were not part of the pool of potential immigrants because their territory quality was already high enough that it could not be improved by relocation. If depletion of potential immigrants had a predominant influence on immigration rate, we would have expected a gradual reduction in immigration rate, with immigration stopping at a range of densities depending on the supply. The observed pattern of a gradual increase in immigration rate with a consistent, abrupt drop to zero only when density was nearly zero suggests that other processes such as avoidance of vacant habitat were involved. Uncertainty about the origin of immigrants is widespread in field studies of density-dependent immigration as is uncertainty about settlement location in studies of density-dependent emigration. This uncertainty creates a risk that studies of density-dependent dispersal may be confounded by correlations between source and settlement densities as result of natural spatial and temporal variation and as a result of unrecognized effects of experimental density manipulations on adjacent populations. Thus, improved insights in future dispersal studies may be gained by monitoring both source and settlement populations, despite the substantial logistical challenges of doing so.

While competition for a variety of resources could explain the negative relationship between density and immigration that we observed over most of the density range, competition for space seems the most likely in our system. In territorial species, space itself can become limiting when all suitable habitat is occupied and territories cannot be compressed below a minimal size [57,70–72]. Immigration then depends on the reoccupation of the territory of a resident that dies, emigrates, or is physically displaced [32,72,73]. In our study, the very low rate of immigration in control sites and the lack of immigration during pre-removal observations on experimental sites follow this pattern. If neighbors of individuals removed at high densities expand their territories into the unoccupied space before it can be discovered by potential immigrants [74], then opportunities for immigration may be limited until densities are low enough that such expansion provides no additional benefit. Mutual defense in which individuals from several territories simultaneously attack a potential immigrant [75] could reduce the probability of both detecting and occupying a vacancy at high densities [76]. In addition, if vacant territories are more likely to be occupied by nearby individuals [77], then removals early in the removal sequence would be more likely to be taken by residents than by immigrants, as operationally defined in our experiment.

Dispersal of post-settlement fish from protected areas to locations where harvesting is allowed, often called ‘spillover’ in the marine fisheries and conservation literature, reduces the protective effect of reserves. It therefore decreases the number of eggs and larvae exported but at the same time provides resources for harvesters outside the reserve [78]. Similar processes have been proposed for terrestrial systems [79]. While spillover can occur through density-independent processes, it has been widely recognized that density-dependent emigration from reserves could enhance spillover [11,80,81]. However, little attention has been given to the potential density-dependence of immigrants settling in harvested areas. Decreasing immigration with decreasing density at low density raises the possibility that excessive harvest outside a reserve might inhibit movement from the reserve. Indeed, conspecific attraction could even induce movement from harvested areas into reserves [11].

In conclusion, this study provides evidence that density can strongly affect immigration in complex ways, with both positive and negative correlations occurring at the same site at different levels of density. While some of the possible mechanisms involved may be limited to very local scales (*e.g.,* depletion of potential immigrants) or be restricted to territorial species (*e.g.,* territorial expansion, mutual defense), others are likely to operate at much larger scales and affect populations with other social systems and dispersal patterns (*e.g.,* resource competition, conspecific attraction, and vacant habitat avoidance). For theoretical explorations of metapopulation dynamics, reserve design and dispersal evolution, our results reinforce recent calls for more recognition of density dependence at all stages of the dispersal process [15], as pioneered by [82]. In addition, our results suggest that simple formulations of dispersal probability as either a difference from or proportion of carrying capacity [83] may be inadequate. It is important that empirical studies of dispersal recognize the potential for density dependence in immigration as well as emigration, cover a broad enough density range to detect such complex patterns, and carefully consider the scaling of immigration rates. Localized harvest and pest control programs offer opportunities to undertake such studies through gradual removal. Despite a long history of using removal experiments to address a variety of questions ranging from habitat preference and social organization to population regulation, interspecific competition, and assemblage structure [57,58,61,84–86] and even emigration [87], ours appears to be the first to use this method to examine immigration over a wide range of densities in the field.

## Acknowledgements

We are indebted to S. Rouleau, A. Ménard, É. Castonguay, J. Grégoire, A. E. Hall, S. Theleme, A. Robillard, V. Duclos, K. Gotanda and B. Spelke for their help in the field. Thanks to T. Avgar, J. Grant, M. Hixon, R. McLaughlin and R. Warner for valuable comments on previous versions of this manuscript. We also want to thank Bellairs Research Institute and High Tide Watersports for their logistical assistance and support during the field work.

## Supporting Information captions

**S1 Figure: Location of experimental and control sites**

Location and names of fringing reefs along the west coast of Barbados, West Indies. Reefs are shown as dark green and the area covered by the Barbados Marine Reserve as light blue. On the left, two reefs illustrate the positions of reef zones, with the spur and groove zone used in this study shown in light green.

**Table: Habitat variables of the seven sites**

Habitat variables of seven sites in which the density of two damselfish species (*Stegastes diencaeus* and *S. adustus*) was manipulated. See Figure 1 for site abbreviations.

**Table: Model support (AICc scores) for per new vacancy scaling**

Per new vacancy scaling. AICc scores (AICc) and Akaike weights (w_*i*_) for each of the six empirical functions predicting observed immigration rate per new vacancy (number of immigrants/number of damselfish removed in a removal event) in relation to population size following each removal event (not including saturation because new vacancies = 0) on seven experimental sites. Sites are ordered as in Figure 1. The mean Akaike weight (Mean w_*i*_) for each function is given at the bottom of each column. For each site and the mean of all sites, the function with the highest support is indicated by bold w_*i*_. See Figure 1 for site abbreviations and Table 1 for the functions.

**Table: Model support (AICc scores) for per capita scaling**

*Per capita* scaling. AICc scores (AICc) and Akaike weights (w_*i*_) for each of the six empirical functions predicting observed *per capita* immigration rate (number of immigrants/number of remaining damselfish) in relation to population size following each removal event (excluding population size = 0) on seven experimental sites. The mean Akaike weight (Mean w_*i*_) for each function is given at the bottom of each column. For each site and the mean of all sites, the function with the highest support is indicated by bold w_*i*_. Sites are ordered as in Figure 1. See Figure 1 for site abbreviations and Table 1 for the functions.

**S2 Figure: Relationships between immigration and the number of damselfish for new vacancy and *per capita* scaling**

Relationships between immigration (per new vacancy scaling: black dots and lines, left ordinate; *per capita* scaling: grey dots and lines, right ordinate) and the number of damselfish remaining following each removal event for each individual site and for all sites combined (upper left panel). Individual sites are ordered by strength of support for a hump-shaped (Ricker) function and abbreviated as in Fig. 1. Lines represent the best fitting empirical function based on maximum likelihood estimates (MLE) of parameter values. For all sites combined, the abscissa indicates the proportion of the initial number of fish on the site before removal. Each point represents the mean ± 1 SE of all points falling within bins of 0.1, with density = 0 shown separately. Sample sizes per bin vary depending on replacement rates. The short inside tick marks on the abscissa represent observed proportion values. The solid lines in the upper left panel represent the significant smoothing term functions from GAMM models and the dashed lines represent the Bayesian credible intervals.

